# An unexpected lack of difference in superoxide/H_2_O_2_ production rates in isolated heart and skeletal muscle mitochondria from a mouse model of Barth Syndrome

**DOI:** 10.1101/2020.05.07.083105

**Authors:** Renata L. S. Goncalves, Michael Schlame, Alexander Bartelt, Martin D. Brand, Gökhan S. Hotamışlıgil

**Author notes:** To whom the correspondence should be addressed: Renata L.S. Goncalves, Sabri Ülker Center for Metabolic Research and Department of Molecular Metabolism, Harvard T.H. Chan School of Public Health, Boston, MA, U.S.A. email: rg.

## Abstract

Barth Syndrome (BTHS) is a rare X-linked genetic disorder caused by mutations in tafazzin and characterized by loss of cardiolipin and severe cardiomyopathy. Mitochondrial superoxide/H_2_O_2_ production has been implicated in the cardiomyopathy observed in different BTHS models. There are at least 11 mitochondrial sites that produce superoxide/H_2_O_2_ at significant rates. Which of these sites generate oxidants at excessive rates in BTHS is unknown. Here, we measured the maximum capacity of superoxide/H_2_O_2_ production from each site in mitochondria isolated from heart and skeletal muscle of the tafazzin knockdown mice (tazkd) at 3, 7 and 12 months of age. Strikingly, the superoxide/H_2_O_2_ production capacities of these sites were overall indistinguishable between tazkd mice and their wildtype littermates across the time points analyzed. The only exception was site G_Q_ in glycerol phosphate dehydrogenase, which was increased in the skeletal muscle of 7 months old tazkd mice. Mitochondrial superoxide/H_2_O_2_ production was also measured *ex vivo* during the oxidation of a complex mixture of substrates mimicking either heart or skeletal muscle cytosol and was found to be indistinguishable between wildtype and tazkd mice. However, we consistently measured decreased FAD-linked respiration in mitochondria isolated from tazkd mice. We conclude that the maximum capacity and *ex vivo* rates of superoxide/H_2_O_2_ production were not increased in mitochondria isolated from heart and skeletal muscle of tazkd mice, despite reduced oxidative capacity. Therefore, it seems unlikely that mitochondrial oxidants contribute to the development of cardiomyopathy in tazkd mice. These observations raise questions about the involvement of mitochondrial oxidants in BTHS pathology.

## Introduction

Cardiolipin (CL) is a unique phospholipid; it contains four acyl chains and is exclusively synthetized and found in the mitochondrial membranes (1). Cardiolipin interacts with a myriad of mitochondrial proteins and this interaction is critical for their function (2, 3). A critical step in cardiolipin synthesis and maturation is the remodeling of its acyl chains to a highly symmetric and unsaturated profile (3, 4). cardiolipin remodeling consists in removing a fatty acyl chain from the nascent molecule, generating a monolysocardiolipin (MLCL), which is further re-acylated by the transacylase, tafazzin (taz) (5) (Figure 1). In the heart and skeletal muscle, taz-mediated cardiolipin remodeling generates tetralinoleoyl cardiolipin (L_4_CL) molecules (4). Upon loss of taz the MLCL/CL ratio is dramatically increased (6, 7). Mutations in the *taz* gene cause a rare X-linked autosomal recessive disease, named after Dr. Peter Barth who first described the syndrome in 1981 (8). Barth syndrome (BTHS) has an early onset, affecting mostly infant male individuals and is characterized by cardiomyopathy, skeletal myopathy, neutropenia, high levels of 3-methylglutaconic acid in the urine and growth delay (3, 8, 9). Nearly a decade ago the first mouse model expressing an inducible short-hairpin RNA to promote taz knockdown (tazkd) became available (6, 10). Tazkd mice recapitulate many aspects of the disease in humans, however the symptoms in the murine model are characterized by a much later onset where cardiomyopathy becomes evident at 8 months (6, 11).

**Figure 1.**
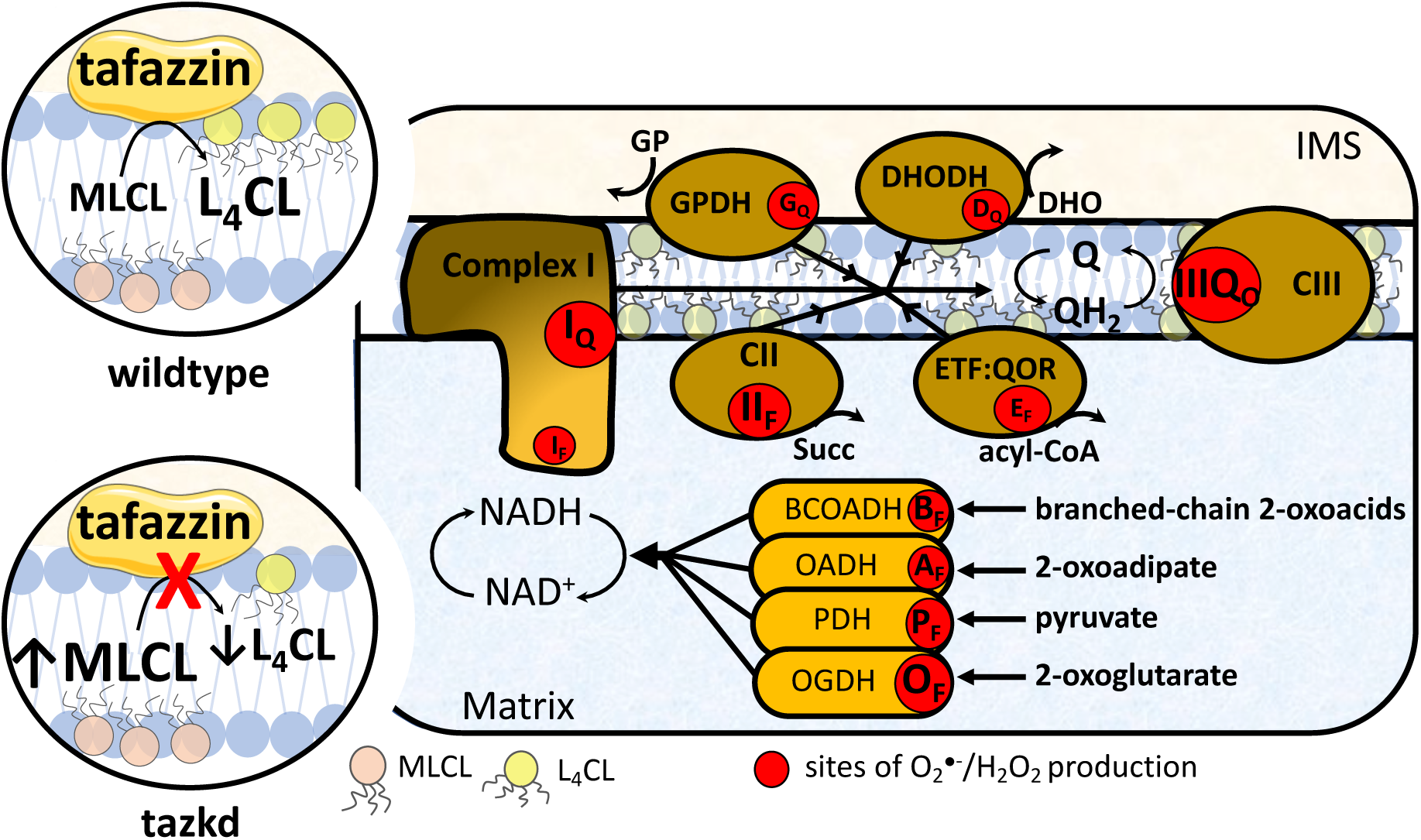
Tafazzin deficiency and the sites of superoxide/H_2_O_2_ in the mitochondria. Tafazzin is an acyltransferase important for remodeling cardiolipin to its physiologically relevant form, tetralinoleoyl cardiolipin (L_4_CL) (circle on top left, wildtype). Mutations in the tafazzin gene alter the cardiolipin profile in the mitochondria and increase the levels of the intermediate, monolysocardiolipin (MLCL) (circle on bottom left, tazkd). Cardiolipin tightly interacts with and stabilize components of the electron transport chain (ETC) and aberrant cardiolipin profile is associated with higher mitochondrial superoxide (O_2_^•-^) and hydrogen peroxide (H_2_O_2_) production. There are eleven sites with the capacity to produce O_2_^•-^/H_2_O_2_ in mitochondria. These sites are associated with the ETC and substrate oxidation enzymes, and are represented by red circles. Reduced substrates from metabolism are transported into the mitochondria where they are oxidized; the electrons enter the ETC via enzymes that operate in close proximity to the NADH/NAD^+^ redox potential and enzymes that operate in close proximity to the ubiquinol/ubiquinone (QH_2_/Q) redox potential. The electrons flow from the NADH- and Q-pool to complex III then to cytochrome *c* and to complex IV, which finally transfers four electrons to oxygen, producing water. The sites associated with the enzymes in the NADH isopotential group and able to generate O_2_^•-^/H_2_O_2_ are the flavin/lipoate of the dehydrogenases of branched chain 2-oxoacids (BCOADH, B_F_), 2-oxoadipate (OADH, A_F_), pyruvate (PDH, P_F_), 2-oxoglutarate (OGDH, O_F_) and the flavin site of complex I (I_F_). The sites in the Q isopotential group are the flavin site of complex II (II_F_) and the electron transfer flavoprotein (ETF) and ETF:ubiquinone oxidoreductase (ETF:QOR) system (E_F_) and the ubiquinone binding sites of dehydrogenases of glycerol phosphate (GPDH, G_F_) and dihydroorotate (DHODH, D_Q_) and the outer quinol site of complex III, III_Qo_. Electrons are transferred from the NADH- to the Q-pool via site I_Q_ in complex I, which has a high capacity for O_2_^•-^/H_2_O_2_ production. The diameters of the red circles are roughly proportional to their mean capacity for O_2_^•-^/H_2_O_2_ generation in heart and skeletal muscle. IMS, intermembrane space; CII, complex II; CIII, complex III.

The abnormal cardiolipin profile due to the loss of taz is associated with mitochondrial dysfunction (3, 9, 10). In particular, mitochondrial production of reactive oxygen species (ROS) – from here referred to as superoxide and H_2_O_2_ – is considered a key process in the pathogenesis of BTHS (3, 9, 12).

Mitochondrial superoxide/H_2_O_2_ generation is not a single process; there are at least eleven sites in the respiratory chain and in different enzymes of substrate oxidation (β-oxidation, tricarboxylic acid cycle, and pyrimidine biosynthesis) that are able to generate significant amounts of these molecules (Figure 1). Each site has its own properties and maximum capacity to produce superoxide/H_2_O_2_ (13). The sites are represented in Figure 1 as red circles and can be didactically separated into two groups based on the redox potential (E*h*) of their redox centers. The sites that operate close to the redox potential of NADH/NAD^+^ (E_h_∼ -280 mV) are the sites in the dehydrogenase complexes of branched-chain 2-oxoacids, 2-oxoadipate, pyruvate and 2-oxoglutarate, sites B_F_, A_F_, P_F_ and O_F_, respectively, and site I_F_ in complex I. These sites belong to the same isopotential group (14–16). The remaining sites, II_F_ in complex II, E_F_ in ETF:QOR G_Q_ and D_Q_ in mitochondrial glycerol 3-phosphate and dihydroorotate dehydrogenase, respectively, and III_Qo_ in complex III, belong to the ubiquinone isopotential group (QH_2_/Q, E_h_ ∼ +20 mV) (14, 17– 21).

The rate of superoxide/H_2_O_2_ production and the contribution from each site heavily depend on the substrates being oxidized (22). For example, in skeletal muscle the rate of superoxide/H_2_O_2_ production during the oxidation of succinate is 5-fold faster than the rate during the oxidation of glutamate plus malate. In addition, the sites engaged during the oxidation of succinate or glutamate plus malate are different. While site I_Q_ is the predominant source of superoxide/H_2_O_2_ when succinate is used as substrate, sites I_F_, III_Qo_ and O_F_ contribute about equally to the rate measured during oxidation of glutamate plus malate (22). There is no *a priori* reason to expect that all sites are equally engaged in superoxide/H_2_O_2_ production in any given context.

*In vivo* or in intact cells, mitochondria metabolize multiple substrates simultaneously, which implies that superoxide/H_2_O_2_ is generated from multiple sites simultaneously. In addition, metabolic effectors, e.g. calcium, ADP or cytosolic pH, which modulate the rate of superoxide/H_2_O_2_ production, vary according to the physiological or pathological context (23–25). Therefore, it is challenging to study site-specific superoxide/H_2_O_2_ generation in intact cells or *in vivo*. Instead, the rate at maximum capacity in isolated mitochondria is widely measured and is often used as a proxy estimation for the generation of these oxidant molecules *in vivo*. However, the maximum capacity cannot be used to predict of the rate of superoxide/H_2_O_2_ production *in vivo* or intact cells (22, 26). To overcome these difficulties, “*ex vivo*” media were carefully designed to mimic the cytosol of skeletal muscle at “rest” and during “exercise”. These media contained the physiological cytosolic concentrations of all substrates and effectors thought to be relevant to oxidative phosphorylation and superoxide/H_2_O_2_ production, allowing more realistic predictions of the rate of superoxide/H_2_O_2_ production from *ex vivo* experiments to *in vivo* conditions (26).

Due to the fact that different groups have reported higher production rates of mitochondrial oxidants in heart and skeletal muscle in BTHS (7, 12, 27– 35), together with the encouraging effects of some antioxidants, especially mitoTEMPO and linoleic acid (12, 35) for improving cardiomyopathy pathogenesis, mitochondrial superoxide/H_2_O_2_ production is considered a key target for therapeutic interventions (3, 11, 12). However, the precise site(s) responsible for the abnormal production of mitochondrial oxidants in different tissues in BTHS are still unknown.

In the present study we isolated heart and skeletal muscle mitochondria from tazkd mice and systematically measured the maximum capacities of superoxide/H_2_O_2_ production from all eleven sites (Figure 1) during the progression of BTSH disease. We took advantage of the late onset of disease in the mouse model to study the kinetics of superoxide/H_2_O_2_ generation in pre-symptomatic 3-month old mice and at later time points where cardiomyopathy and skeletal myopathology are already established (7- and 12-month-old mice). Unexpectedly, our results indicate that there is no difference in the rate of superoxide/H_2_O_2_ production between tazkd mice and their wildtype littermates in heart or skeletal muscle mitochondria during the course of the disease progression.

To estimate the production rate of mitochondrial oxidants *in vivo* in these tissues, superoxide/H_2_O_2_ was measured “*ex vivo*” in media designed to mimic the cytosol of skeletal muscle (26) and heart. Strikingly, “*ex vivo*” rates of superoxide/H_2_O_2_ production were not altered in tazkd mice in mitochondria from either tissue despite the consistent decrease in oxygen consumption and *in vivo* exercise intolerance. In this study, we provide a broad and systematic characterization of the maximum capacities of sites of superoxide/H_2_O_2_ production in heart and muscle mitochondria from tazkd mice during the progression of BTHS. Our data suggest that i) the involvement of mitochondrial superoxide/H_2_O_2_ production in BTSH pathogenesis should be interpreted carefully and/or ii) in tazkd mice, cardiac and muscle pathologies may not be related to the production of mitochondrial oxidants.

## Results

### Tafazzin-deficiency, cardiolipin levels and metabolic characteristics

The tazkd mouse was the first animal model created to study how loss of tafazzin induces Barth syndrome cardiomyopathy (6, 10). In this model, an shRNA expression system under the control of doxycycline decreases tafazzin mRNA and protein levels (6, 10). In our experiments, transgenic males (B6.Cg-*Gt(ROSA)26Sor*^*tm37(H1/tetO-RNAi:Taz)Arte*^/ZkhuJ) were crossed with wildtype females and kept on standard chow diet containing doxycycline. Tafazzin knockdown was induced prenatally and male littermates (wildtype and tazkd) were kept on doxycycline-containing rodent chow throughout the experiment. Figures 2A and B show that tafazzin mRNA levels in the tazkd mice were <30% of their wildtype littermates in both heart and skeletal muscle. The decreased tafazzin mRNA levels were accompanied by a more dramatic decrease in tafazzin protein levels, reaching undetectable levels in the heart as early as 3 months of age and up to one year old (Figure 2C, heart). In the tazkd mice skeletal muscle tafazzin protein levels were <15% of wildtype mice (Figure 2C, skeletal muscle 12 months old). To confirm that lower tafazzin levels resulted in altered cardiolipin remodeling, which is a characteristic of heart and skeletal muscle from BTHS patients, we measured the content of cardiolipin and monolysocardiolipin. In tazkd mice, MLCL accumulated in both tissues, which resulted in significant increases in the MLCL/CL ratio (Figure 2D) (6).

**Figure 2.**
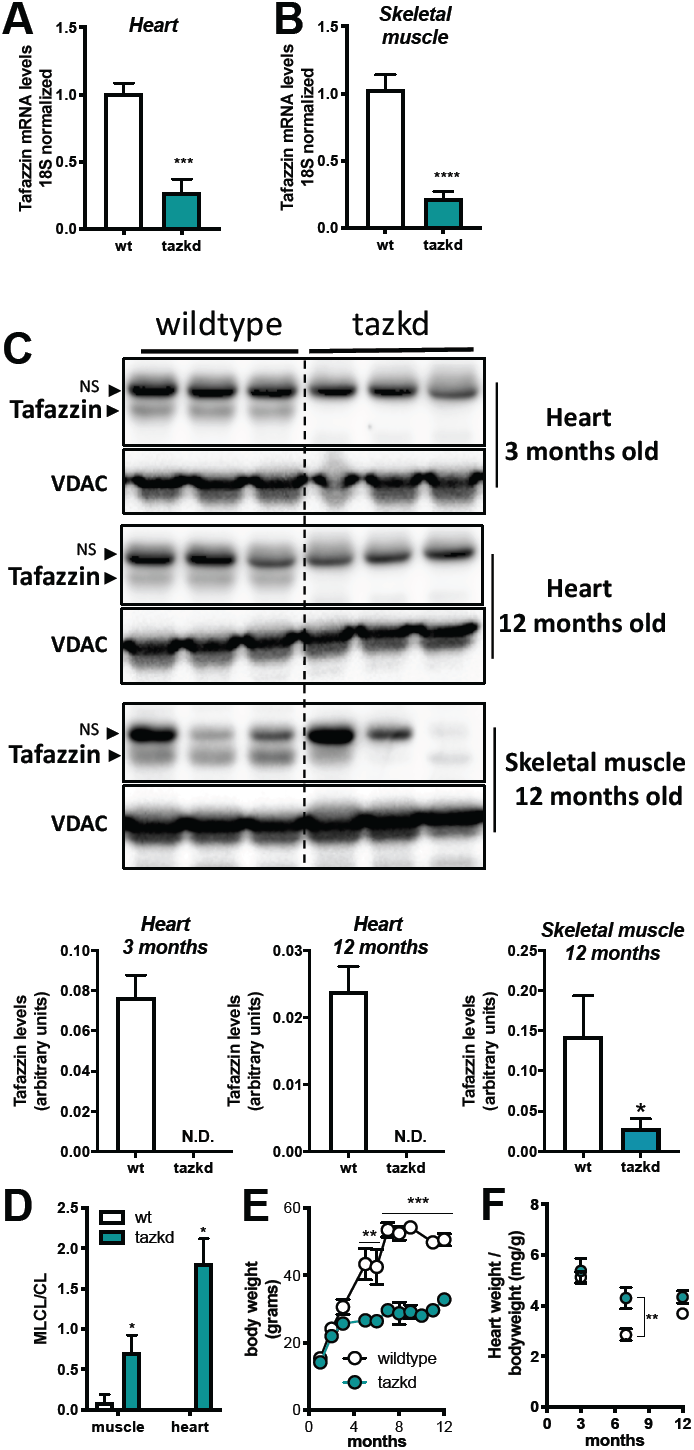
Tafazzin knockdown results in altered cardiolipin levels and protects from weight gain. **A**. Tafazzin mRNA levels in the heart and **B**. in the skeletal muscle of wildtype and tafazzin knockdown mice (tazkd). **C**. Upper, Western blot of tafazzin protein in heart extracts of 3 and 12 month-old wildtype and tazkd mice and in skeletal muscle extracts of 12 month-old wildtype and tafazzin knockdown mice probed with anti-tafazzin antibodies (61). VDAC was used as a loading control. Bottom, densitometric analysis of Western blots from the upper panel C, tafazzin levels were normalized by VDAC levels, NS, non-specific band. **D**. Monolysocardiolipin (MLCL) and cardiolipin (CL) levels were determined by high-performance liquid chromatography in heart and skeletal muscle isolated mitochondria from wildtype and tazkd mice; an elevated ratio of MLCL/CL indicates a defect in CL remodeling. **E**. Body weights of wildtype and tazkd mice. **F**. Heart weight normalized by total body weight. Values are mean ± SEM, n≧3. *, p<0.05; ***, p<0.001 using Students t-test. **, 2-way ANOVA, Sidak’s post-test.

Consistent with previous reports, wildtype mice on doxycycline chow gained significantly more weight than their tazkd littermates (Figure 1E) (30, 33). Tazkd fat (FM) and fat free/lean (FFM) mass were ∼3.5- and 1.4-fold lower than wildtype mice, respectively (Supplementary Figure 1A). We used indirect calorimetry to better understand whether differences in food intake or energy expenditure were responsible for the alteration in bodyweight and composition between wildtype and tazkd mice 6-7 months of age. There was no difference in locomotor activity (supplementary Figure 2B). Food intake was higher in tazkd than wildtype mice (Supplementary Figure 2C), which was paralleled with a higher respiratory exchange ratio (RER) in tazkd mice (Supplementary Figure 2D and E). The RER is the ratio between the volume of CO_2_ produced to the volume of O_2_ consumed (VCO_2_/VO_2_) and indicates the metabolic substrate preferentially oxidized by the mouse. With an RER closer to 1.0 tazkd mice may oxidize exclusively carbohydrates in contrast to wildtype mice, which may have more substrate oxidation flexibility (RER = 0.9). Energy expenditure (EE) was also significantly higher in tazkd mice when normalized by total body weight (supplementary Figure 2 F-G). However, when EE was normalized by the metabolically active lean mass instead, the difference was no longer observed (Supplementary Figure 2H-I). These results indicate that the differences in EE between 6-7 months old wildtype and tazkd are mostly explained by the differences in body weight rather than differences in genotype. However, it is possible that tazkd have slightly higher EE, as we found that at 3 months of age, when body weight was not yet different, EE was already higher in tazkd mice compared to wildtype controls (Supplementary Figure 2 J-K), which is also in line with previous reports (30). Heart weight normalized by body weight was 34% higher in tazkd mice at 7 months of age indicating the characteristic heart hypertrophy (Figure 2F) (33).

### Tafazzin-deficiency impairs mitochondrial function and endurance capacity

Mitochondrial dysfunction, exercise intolerance, and decreased muscle strength and contractility are some of the hallmarks of cardioskeletal myopathy in BTHS (6, 10, 36, 37). Since cardiolipin interacts with every component of the electron transport chain, it is not surprising that in tazkd mice substrate oxidation and ATP synthesis are compromised (9, 38, 39).

Due to the late onset of the disease in this mouse model (6) and previous reports that at early age mitochondrial oxygen consumption in cardiac mitochondria is not altered (40), we isolated mitochondria from 12-month old mice. As shown in Figure 3 A-B, loss of mature cardiolipin in tazkd mice resulted in lower ADP-stimulated oxygen consumption in heart and skeletal muscle isolated mitochondria oxidizing succinate + rotenone as substrate. Powers et al. (36), previously reported that 4-5-month old tazkd and their wildtype littermates can run for about 36 min to a total of

**Figure 3.**
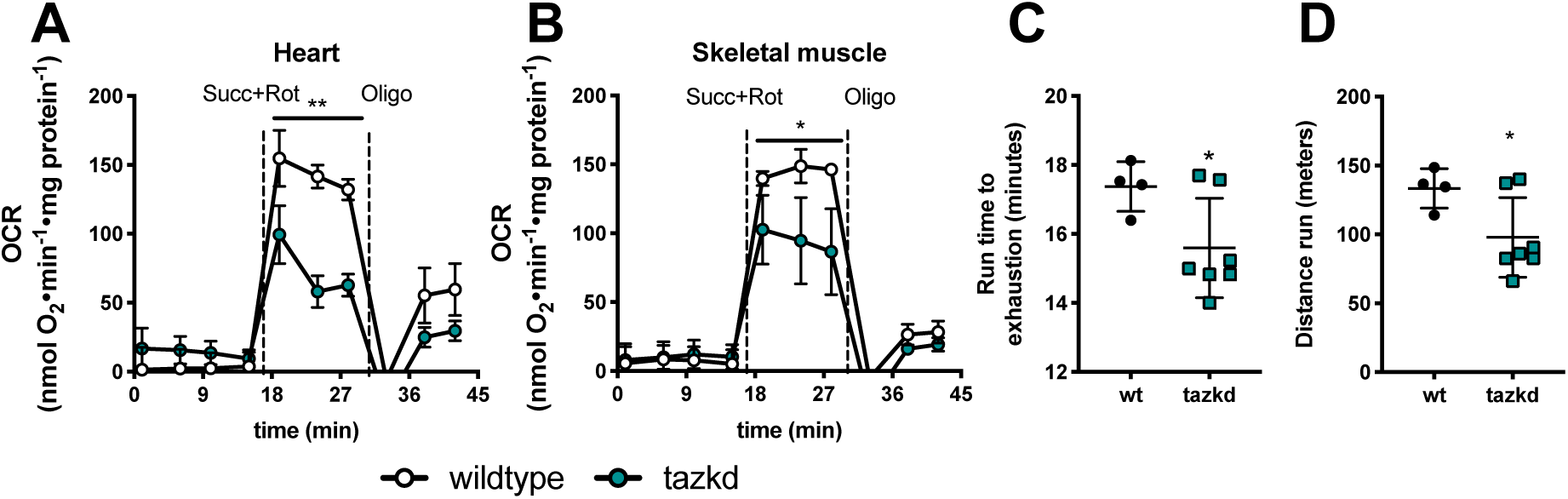
Mitochondrial oxygen consumption rates (OCR) and animal endurance capacity in 12 month-old tazkd mice. **A**. Oxygen consumption rates of mitochondria isolated from heart and **B**. skeletal muscle oxidizing 5 mM succinate in the presence of 4 µM rotenone. Basal rates were measured in the presence of 1 mM ADP. **C**. Duration and **D**. distance run by 12 month-old wildtype and tazkd mice until exhaustion. Values are mean ± SEM, n=3. *, p<0.05 using Students t-test.

507.4 m. When we challenged 12-month old tazkd and wildtype mice in a treadmill, exercise endurance, as measured by running time and distance run, was significantly lower in tazkd mice (Figure 3 C-D). Although, the endurance difference between the genotypes was small, at 1 year of age wildtype mice were obese and weighed 1.6-fold more than tazkd mice (51 ± 0.9 g, n=11 vs 32.5 ± 0.9 g, n=17; p<0.0001). Considering that the endurance of obese mice is 5-6-fold lower than their lean littermates (41), the results from Figure 3 C-D stress the high degree of exercise intolerance of tazkd mice.

### Maximum capacities of the eleven sites of superoxide/H_2_O_2_ production

Loss of cardiolipin in BTHS is associated with increased production of mitochondrial oxidants (7, 12, 27–35), which have been considered to have a causative role in the development of cardiomyopathy (12, 34, 35). However, other reports have failed to find differences in mitochondrial oxidant production in cardiac tissue of tazkd mice (40, 42). There are eleven sites associated with the electron transport chain and tricarboxylic acid cycle that are known to produce superoxide/H_2_O_2_ at significant rates. Their maximum capacities and native uninhibited rates of superoxide/H_2_O_2_ have been systematically characterized in rat skeletal muscle mitochondria (13, 26). Complex I and complex III are often credited as the main source of mitochondrial oxidants (43–45). Indeed, complex I-III activity is decreased in tazkd cardiac mitochondria (36, 37, 42), however, whether this reflects higher superoxide/H_2_O_2_ production from these sites or other adjacent sites in tazkd mice is not yet known. We hypothesized that mitochondrial superoxide/ H_2_O_2_ generation would be higher in tazkd mice cardiac and skeletal muscle mitochondria and our objective was to identify the site(s) that produced excess oxidants.

#### Cardiac mitochondri

To gain insight into the dynamics of superoxide/H_2_O_2_ production in cardiac mitochondria in BTHS, the maximum capacities of the eleven sites were measured in heart mitochondria isolated from wildtype and tazkd mice at 3, 7 and 12 months of age (Figure 4). To compare the maximum rate of H_2_O_2_ generation from each site between the two genotypes, we pharmacologically isolated the sites in intact mitochondria by providing the appropriate combination of inhibitors and substrates at saturating concentrations (see experimental procedures). The sites with the highest maximum capacities were: III_Qo_ in complex III, I_Q_ in complex I and II_F_ in complex II (Figure 4). The maximum capacities of the sites I_Q_ and III_Qo_ were higher at 7 months of age in wildtype mice (2-way ANOVA, age: p<0.003, interaction p=0.1727) but this difference was absent in tazkd mice (2-way ANOVA, age: p=0.6422, interaction: p=0.9998).

**Figure 4.**
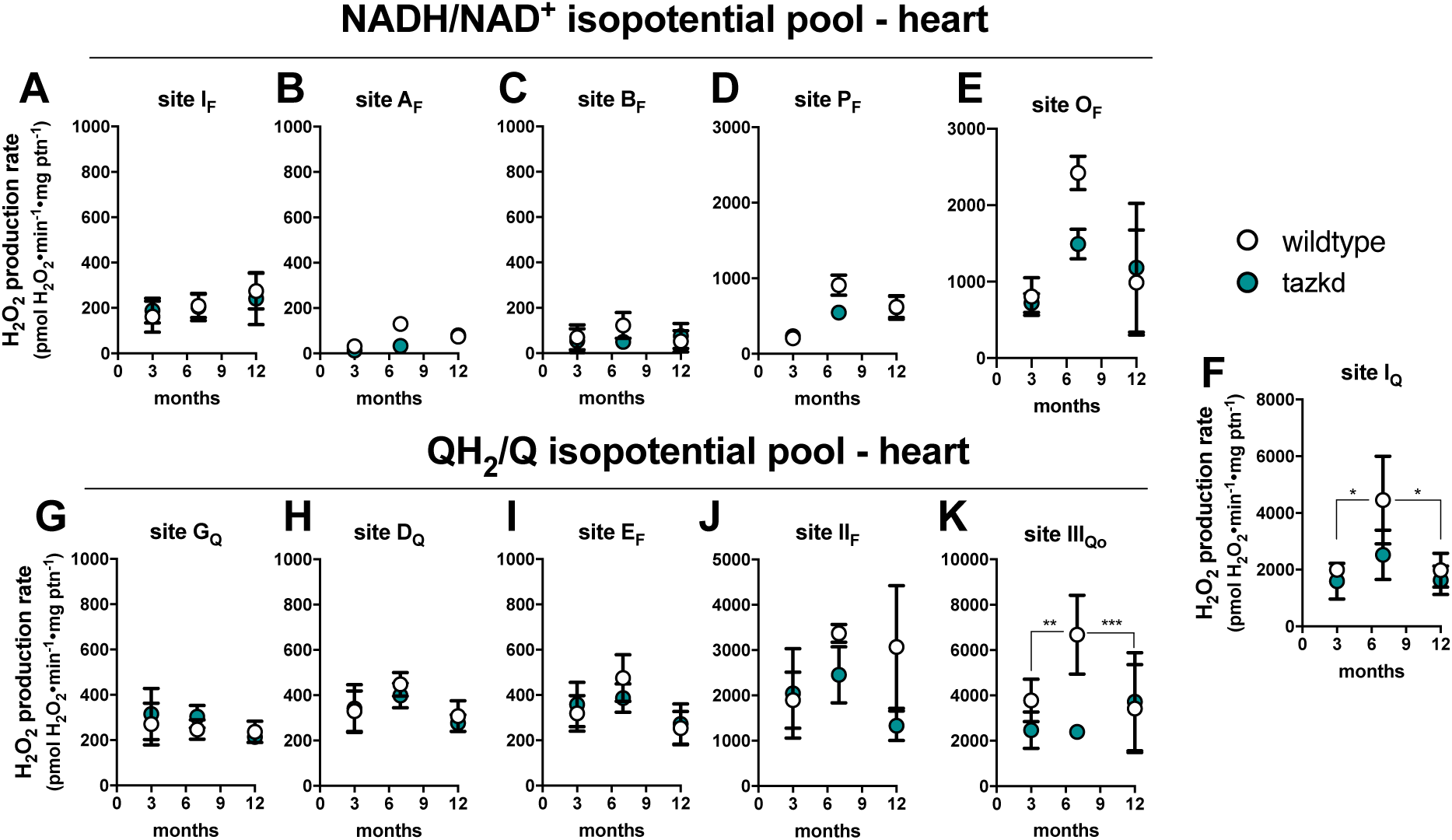
Maximum rate of superoxide/hydrogen peroxide production in isolated heart mitochondria from wildtype and tazkd mice at 3, 8 and 12 months of age. The rate of superoxide/hydrogen peroxide generated by sites associated with the NADH/NAD^+^ (**A-E**) and QH_2_/Q (**G-H**) isopotential pools. In the NADH/NAD^+^ isopotential pool, the rate of superoxide/H_2_O_2_ production was measured from the flavin (F) binding sites of: (**A**) complex I (site I_F_), (**B**) 2-oxoadipate dehydrogenase (site A_F_), (**C**) branched chain 2-oxoacid dehydrogenase (site B_F_), (**D**) pyruvate dehydrogenase (site P_F_), and (**E**) 2-oxoglutarate dehydrogenase (site O_F_). Complex I produces superoxide/H_2_O_2_ from two sites: (**A**) site I_F_ and (**F**) site I_Q_ (ubiquinone (Q) binding site). In sites associated with the QH_2_/Q isopotential pool, the rate of superoxide/hydrogen peroxide production was measured from (**G**) site G_Q_ (glycerol 3-phosphate dehydrogenase), (**H**) site D_Q_, in dihydroorotate dehydrogenase, (**I**) site E_F_, in the electron transfer flavoprotein (ETF) and ETF:ubiquinone oxidoreductase (ETF:QOR) system, (**J**) site II_F_, in complex II, and (**K**) site III_Qo_, in complex III. Values are mean ± SEM n≧3 and were normalized by mitochondrial protein (ptn). 2-way ANOVA was used to determine significance. H_2_O_2_ production rate from sites I_Q_ and III_Qo_ is greater in 7 month-old compared to 3 and 12 month-old in wildtype (2way ANOVA, age: p<0.003, interaction p=0.1727) but not in tazkd mice (2-way ANOVA, age: p=0.6422, interaction: p=0.9998).

Figure 4 shows the rates of H_2_O_2_ generation from the eleven sites grouped according to their operating redox potentials. Sites A_F_, B_F_, P_F_ and O_F_ in the matrix dehydrogenases of 2-oxoadipate, branched chain amino acids, pyruvate and 2-oxoglutarate, respectively, belong to the NADH/NAD^+^ isopotential pool, together with site I_F_ in complex I (Figure 4 A-E). The electrons from NADH, generated by these dehydrogenases, are transferred via site I_F_ to site I_Q_ in complex I (Figure 1F), and then to the next isopotential group, at the redox potential of QH_2_/Q (Figure 4 G-K). In this isopotential group are sites G_Q_, D_Q_, E_F_, II_F_, which reduce ubiquinone (Q) to ubiquinol (QH_2_), and the site in complex III (site III_Qo_) that oxidizes the ubiquinol and ultimately transfers the electrons to cytochrome *c*, complex IV and O_2_.

Strikingly, the rate of superoxide/H_2_O_2_ production was not increased in tazkd mice at any time point analyzed compared to the matched wildtype controls (2-way ANOVA was used to identify significant associations with age and genotype for each site). Instead, there was a trend in the opposite direction, and the rate of H_2_O_2_ production from sites O_F_ and III_Qo_ were actually lower in tazkd compared to wildtype control mice at 7 months of age (Figure 4 E and K) (site O_F_ 2423 ± 219 vs 1491 ± 195; site III_Qo_ 6680 ± 1735 vs 2390 ± 172.5, mean ± SEM pmol H_2_O_2_•min^-1^•mg protein^-1^; n = 3).

#### Skeletal muscle mitochondria

The rate of superoxide/H_2_O_2_ production from the eleven distinct sites were also measured in mitochondria isolated from skeletal muscle of wildtype and tazkd mice at 7 and 12 months of age (Figure 5) using the same approach described above. The sites with the highest maximum capacities were: site III_Qo_, site I_Q_, site II_F_ and site O_F_ (in the 2-oxoglutarate dehydrogenase complex) (Figure 5). The rate of H_2_O_2_ production from site III_Qo_ was higher in wildtype and tazkd mice at 7 months of age (2-way ANOVA p<0.001, Sidak’s post-test). In tazkd mice, H_2_O_2_ production from site G_Q_, in glycerol phosphate dehydrogenase (GPDH), was the only site to be significantly higher when compared to the wildtype littermates (1215 ± 34, n=3 vs 1802 ± 100, n=3, p=0.0011; 2-way ANOVA, Sidak’s post-test followed by Bonferroni correction for testing multiple sites).

**Figure 5.**
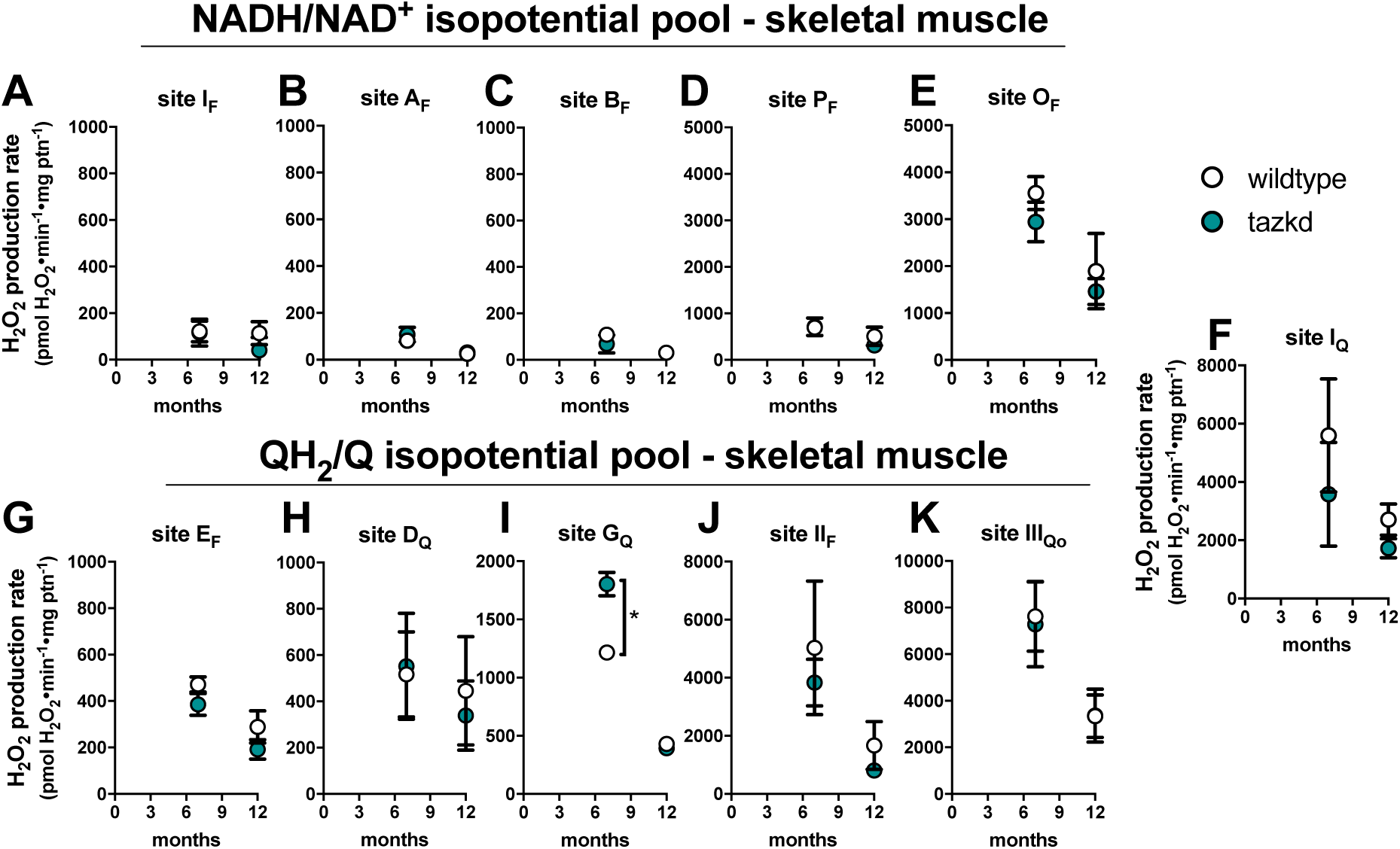
Maximum rate of sites generating superoxide/hydrogen peroxide in isolated skeletal muscle mitochondria from wildtype and tazkd mice at 3, 8 and 12 months of age. The rate of superoxide/hydrogen peroxide generated by sites associated with the NADH/NAD^+^ (**A-E**) and QH_2_/Q (**G-H**) isopotential pools. In the NADH/NAD^+^ isopotential pool, the rate of superoxide/H_2_O_2_ production was measured from the flavin (F) binding sites of: (**A**) complex I (site I_F_), (**B**) 2-oxoadipate dehydrogenase (site A_F_), (**C**) branched chain 2-oxoacid dehydrogenase (site B_F_), (**D**) pyruvate dehydrogenase (site P_F_), and (**E**) 2-oxoglutarate dehydrogenase (site O_F_). Complex I produce superoxide/H_2_O_2_ from two sites: (**A**) site I_F_ and (**F**) site I_Q_ (ubiquinone (Q) binding site). In sites associated with the QH_2_/Q isopotential pool, the rate of superoxide/hydrogen peroxide production was measured from (**G**) site E_F_, in the electron transfer flavoprotein (ETF) and ETF:ubiquinone oxidoreductase (ETF:QOR) system, (**H**) site D_Q_, in dihydroorotate dehydrogenase, (**I**) site G_Q_ (glycerol 3-phosphate dehydrogenase), (**J**) site II_F_, in complex II, and (**K**) site III_Qo_, in complex III. Values are mean ± SEM, n≧3 and were normalized by mitochondrial protein (ptn). *,p<0.001 2-way ANOVA.

The assessment of H_2_O_2_ production rate at maximum capacity fundamentally measures the V_max_ of electron leak to O_2_ and is broadly used to report mitochondrial oxidant production. The advantages of assessing H_2_O_2_ production at maximum capacity are discussed elsewhere (13). However, these measurements are non-physiological, as conventional substrates are presented in excess together with respiratory chain inhibitors. Under native conditions (i.e. in the absence of inhibitors) the rate of H_2_O_2_ generation is much lower (14, 22, 26), therefore maximum capacities cannot be used to predict the actual rate of H_2_O_2_ generation in intact cells or *in vivo*.

#### *Ex vivo* rates of mitochondrial H_2_O_2_ production at “rest”

In intact cells and tissues multiple substrates are oxidized simultaneously and various sites generate superoxide/H_2_O_2_ simultaneously and at different rates (26, 46). The rate of H_2_O_2_ production *in vivo* is tissue-specific and depends on many factors, including the substrates being preferentially oxidized and the abundance of the protein complex that contains the site from which the electrons leak to oxygen generating superoxide and then H_2_O_2_. For example, the rate of superoxide/H_2_O_2_ production from site G_Q_ in 7-month old mice is ∼5-fold higher in mitochondria from skeletal muscle than in those from heart (Figure 4G and 5I), which is correlated to the level of GPDH in these tissues (20). To approach physiology and assess the rate of superoxide/H_2_O_2_ in isolated mitochondria under more realistic conditions, we carefully designed media that mimicked the cytosol of skeletal muscle at rest (not contracting) (26) and heart at rest (contracting but not under unusual load). These media contained all the relevant substrates and effectors considered relevant to mitochondrial electron transport and superoxide/H_2_O_2_ production at the physiological concentration found in the cytosol of heart and skeletal muscle at “rest” [see supplementary table 1 and (26)]. We refer to the rates of superoxide/H_2_O_2_ obtained in these media as “native” “*ex vivo*” rates. In Figure 6 we used these experimental systems to determine the rates of mitochondrial superoxide/H_2_O_2_ production in conditions mimicking cardiac and skeletal muscle at rest.

**Figure 6.**
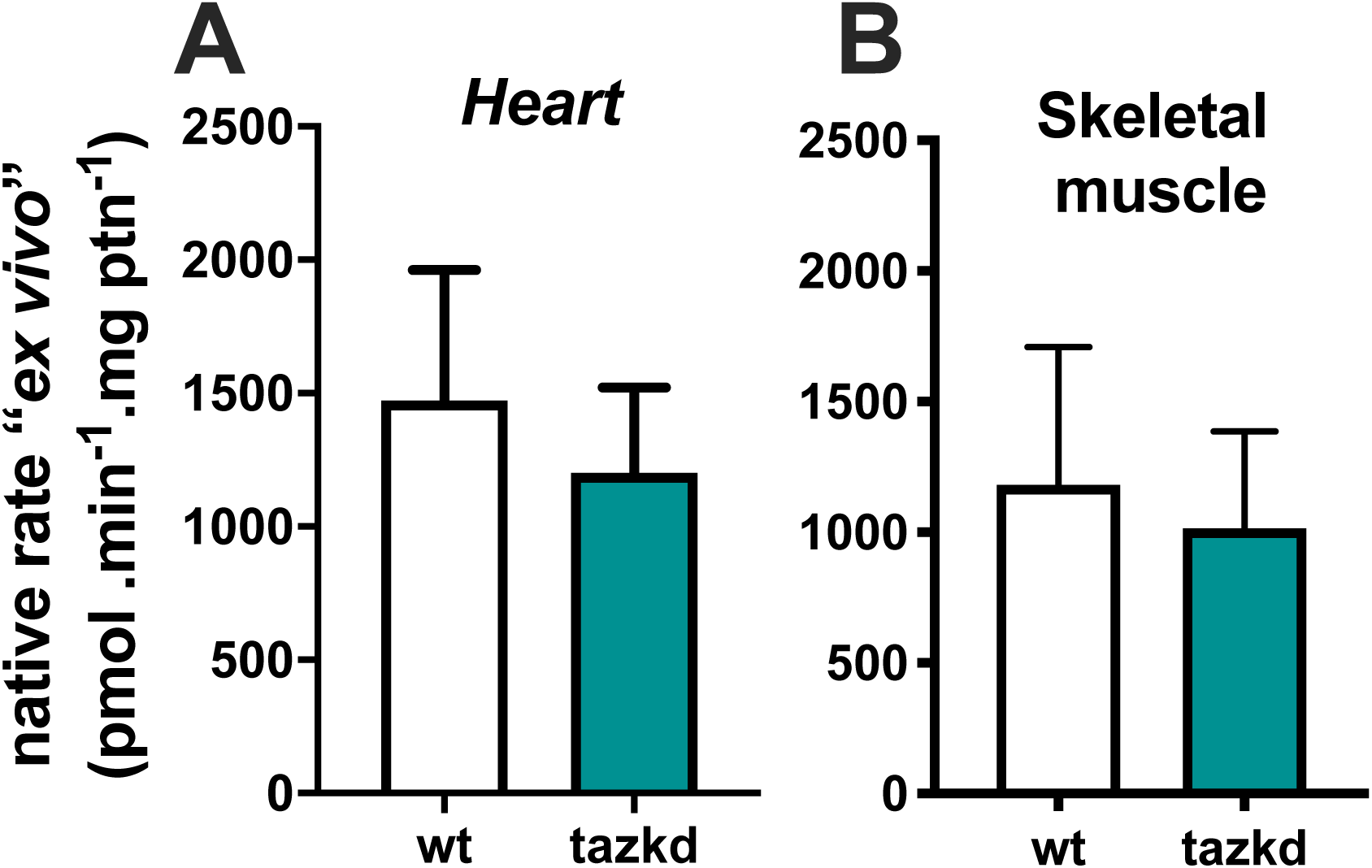
Native *ex vivo* rates of superoxide/hydrogen peroxide production from isolated mitochondria from 12 month-old mice. Mitochondrial H_2_O_2_ production rate was measured in complex media mimicking the cytosol of (A) heart and (B) skeletal muscle “at rest” (see methods for media composition). Values are mean ± SEM, n≧3 normalized by mitochondrial protein (ptn). p>0.05 Student’s t-test.

The rest state in our system was defined by a low rate of ATP synthesis, which was achieved by the addition of oligomycin. Although not ideal, oligomycin is necessary due to the contaminating ATPases present in the mitochondrial preparation.

Mitochondria isolated from heart and skeletal muscle from 12-month old mice were incubated in their respective media. Strikingly, the “*ex vivo*” rates of superoxide/H_2_O_2_ production by mitochondria from wildtype and tazkd mice were indistinguishable when using mitochondria from heart (Figure 6A) or from skeletal muscle (Figure 6B).

Taken together, we detected no difference in the rates of superoxide/H_2_O_2_ production from most sites in heart or skeletal muscle mitochondria isolated from tazkd mice compared to wildtype controls. The only exception was the capacity of site G_Q_, which was significantly increased in the skeletal muscle from tazkd mice at 7 months of age (Figure 5I).

## Discussion

In any intact system, whether cells or tissues *in vivo*, the rate of mitochondrial oxidant species production is the sum of superoxide/H_2_O_2_ production from up to eleven sites (Figure 1). Mitochondrial oxidants play an important role in the development of many pathological states. In BTHS, excess mitochondrial oxidants are thought to have a causative role in the development of the cardioskeletal myopathy. However, these molecules are also important for normal cell signaling and physiology (45), which may explain why supplementation with broad and unselective antioxidants to prevent BTHS cardioskeletal myopathy failed (11). In addition, such non-specific antioxidants may not target the appropriate oxidant species in the right cellular compartment. Still, a major gap in the field is the ability to identify and selectively target the site or sites from which the excess superoxide/H_2_O_2_ arises while at the same time preserving physiological oxidant production from other sites. We hypothesized that in tazkd mice, mitochondrial superoxide/H_2_O_2_ production is non-homogeneous and specific sites (perhaps those in the respiratory chain complexes directly affected by cardiolipin) may generate mitochondrial oxidants at abnormally high rates. The excess superoxide/H_2_O_2_ production could be then normalized using a new generation of suppressors of mitochondrial electron leak at sites I_Q_ and III_Qo_, S1QELs and S3QELs (47, 48).

The present study set out i) to identify which site(s) have increased capacity to generate excess superoxide/H_2_O_2_ in heart and skeletal muscle of tazkd mice and ii) to measure the physiologically-relevant rates of superoxide/H_2_O_2_ production from these sites *“ex vivo”* using complex media mimicking cardiac and skeletal muscle cytosol at rest. Strikingly, we found that the overall maximum capacities of superoxide/H_2_O_2_ generation in mitochondria isolated from cardiac and skeletal muscle were not different between the genotypes. Site G_Q_ was the only exception and had a higher capacity in tazkd mice than controls at 7 months of age. In the more physiological approach we found that there was no difference in the “*ex vivo*” rates of superoxide/H_2_O_2_ production in mitochondria from heart and skeletal muscle between tazkd mice and their littermate wildtype controls.

Mitochondrial ATP supply is crucial to satisfy the high energetic demand of cardiac and skeletal muscle. Therefore, it is not surprising that altered cardiolipin composition in BTHS results in cardioskeletal myopathy. Loss of cardiolipin alters mitochondrial morphology (49) and causes the destabilization of the components of the oxidative phosphorylation (38, 39, 50). Tafazzin deficiency in BTHS has been overwhelmingly reported to be associated with increased production of mitochondrial oxidants (7, 12, 27–35). Indeed, targeting mitochondrial oxidants with mitoTempo and linoleic acid ameliorated myocardial dysfunction in models of BTSH (12, 35), suggesting that mitochondrial oxidants have a causative role in BTHS pathogenesis.

The reliable measurement of mitochondrial superoxide/H_2_O_2_ levels and production rates in tissues *in vivo* and intact cells is very challenging due to the lack of specific probes. For example, dichlorodihydrofluorescein (DCFH_2_), which has been widely used to report differences in oxidant production in BTHS (31, 32, 34, 49), does not directly react with H_2_O_2_ (51). Importantly, H_2_O_2_-mediated DCFH_2_ oxidation is dependent on iron uptake (52) and loss of cardiolipin increases levels of iron uptake genes (53). Importantly, DCFH_2_ itself can generate superoxide in the presence of oxygen (51). MitoSOX has also been widely used to measure levels of mitochondrial superoxide in intact cells and has been used to report mitochondrial oxidants in different BTHS models (7, 12, 29, 34, 35). However, the results obtained with mitoSOX should be interpreted with caution (51). MitoSOX red fluorescence can be the result of unspecific oxidation in the presence of iron and therefore does not solely or even dominantly reflect superoxide levels. Also, its accumulation in the mitochondria is dependent on both mitochondrial and plasma membrane potentials and it is advised for the signal to be normalized to correct for this effect. Lastly, depending on the concentrations, mitoSOX disturbs the respiratory chain, which will impact superoxide generation (54). Taken together, the interpretation of data obtained using these probes should be reevaluated. Finally, the other probe commonly used to report H_2_O_2_ in isolated mitochondria in BTHS is Amplex ultraRed (27, 30, 33, 40). Although one should be aware that this probe can also provide some unspecific signal depending on the source of mitochondria (51, 55), Amplex ultraRed is more specific for H_2_O_2_ and is more reliable than its predecessor, Amplex Red (56). Interestingly, the published data using Amplex ultraRed is more heterogeneous as some studies show a higher rate of H_2_O_2_ generation in the context of tazkd (30, 33) while others report no difference (33, 40).

In the present work we provide a comprehensive assessment of the rate of superoxide/H_2_O_2_ generation from each individual site in heart and skeletal muscle mitochondria during the progression BTHS in tazkd mice. Unexpectedly, we found no major differences in the maximum capacities of the different sites (Figures 4 and 5) despite a clear phenotype of reduced oxidative capacity (Figure 2 and Supplementary Figure 1). This unexpected lack of association between mitochondrial superoxide/H_2_O_2_ production and the cardioskeletal myopathology in tazkd mice was also observed in MCAT-tazkd mice (33). In that study, tazkd mice were crossed with a mouse line expressing the H_2_O_2_-degrading enzyme catalase (CAT) targeted specifically to the mitochondrial matrix. In these MCAT-tazkd mice, mitochondrial matrix H_2_O_2_ levels in the heart were lower but this was insufficient to improve the cardioskeletal myopathologies (33). Similarly, it was recently reported that the mitochondrial-targeted antioxidant mitoQ and the general antioxidant n-acetylcysteine did not improve cardiopathologies in tazkd mice^1^.

To overcome the drawbacks of trying to measure levels and rates of production of mitochondrial oxidants in intact cells with DCFH_2_ and mitoSOX and to be able to approach physiology using isolated mitochondria, we designed “*ex vivo*” media that mimicked the cytosol of heart and skeletal muscle at rest (Figure 6, Supplementary Table 1). This approach was previously validated in three different media designed to mimic the cytosol of skeletal muscle at rest and during mild and intense exercise (26). The rate of superoxide/H_2_O_2_ generation was higher at “rest” than in conditions mimicking exercise. At “rest” sites I_Q_ and II_F_ accounted for half of the total measured rate of superoxide/H_2_O_2_ production. Interestingly, the contribution of site III_Qo_, which has a high capacity for generating superoxide/H_2_O_2_, was similar to the low capacity site I_F_ (26). The contributions of sites I_Q_ and III_Qo_ were further confirmed in C2C12 myoblasts using S1QELs and S3QELs (57). From these results we concluded that the mitochondrial rate of superoxide/H_2_O_2_ production measured at maximum capacity (where individual substrates are present in excess in the presence of inhibitors) should not be used to predict the rates in tissues *in vivo*.

Although the more complex “*ex vivo*” approach offers notable advantages, because it uses all the relevant metabolites and effectors at the physiological concentrations found in the cytosol of cardiac and skeletal muscle cells (Supplementary Table 1), it is still not perfect. The media design for these experiments were based on the levels of metabolites and effectors found in the cytosol of rat skeletal muscle at rest and cardiac muscle cells in beating hearts not under load [Supplementary Table 1 and (26)] and may not be a simulacrum of the metabolites in the mouse tissues, particularly since oligomycin was used to inhibit the mitochondrial ATP synthase and prevent high recycling of ATP produced by contaminating ATPases. In addition, and perhaps critically, the concentrations of these metabolites are unknown in tazkd cardiac and skeletal cells, so they were assumed to be similar between wildtype and tazkd mice.

Finally, the doxycycline diet, necessary to induce the short-hairpin RNA against taz, promoted weight gain in wildtype mice (Figure 1 and Supplementary Figure 1) (6, 30). In our facility, 8 months old wildtype mice weighed as much as genetically obese, *ob/ob*, mice (41). Obesity is associated with altered glucose homeostasis and insulin resistance, which has been previously reported in tazkd wildtype littermate mice on the doxycycline diet (30). Mitochondrial generation of superoxide/H_2_O_2_ is higher in the heart and skeletal muscle of different models of obesity (58, 59). Therefore, from 8-12 months of age, we cannot rule out the possibility that increased mitochondrial oxidant production in the heart and skeletal muscle in the tazkd could be masked by an independent effect of obesity and associated changes on mitochondrial oxidant production in the wildtype controls. It is important to note, however, that at 3-months of age, there was no difference in body weight between genotypes and therefore, the potential confounding effect of body weight was absent. Under these conditions, we still failed to detect any differences in the capacity of superoxide/H_2_O_2_ production between wildtype and tazkd (Figure 4).

## Conclusion

We used a systematic approach to determine the maximum capacities and physiologically relevant rates of mitochondrial superoxide/H_2_O_2_ production from all known sites associated with substrate oxidation in mitochondria isolated from tazkd mice and their wildtype littermates. Our results strongly support the nascent idea that cardioskeletal myopathy in tazkd is not associated with increased production of mitochondrial oxidants. Future studies using other models of BTHS, such as the tafazzin knockout mice, will be important to elucidate the relationship between mitochondrial oxidant generation and BTSH cardioskeletal pathologies.

## Experimental Procedures

### Animals, Mitochondria and Reagents

All animal procedures presented here were approved by Institutional Animal Care and Use Committee (IACUC) of Harvard University. Mice were kept on 12-hour light/12-hour dark cycle in the Harvard T.H. Chan School of Public Health pathogen-free barrier facility. Tafazzin knockdown male mice were obtained from Jackson laboratory (B6.Cg-*Gt(ROSA)26Sor*^*tm37(H1/tetO-RNAi:Taz)Arte*^/ZkhuJ, stock #014648) and crossed with C57BL/6NJ females. Pregnant females were provided with 625 mg/kg doxycycline-containing chow (C13510i, Research Diets) to induce tafazzin silencing in the pups during early development. After weaning, wildtype and heterozygous male mice were kept on doxycycline chow for the remainder of the study. All mice were checked to be negative for nicotinamide nucleotide transhydrogenase (nnt). Skeletal muscle mitochondria from 1-2 wildtype and tazkd mice were isolated from hind limb at 4°C in Chappell-Perry buffer [CP1-0.1 M KCl, 50 mM Tris, 2 mM EGTA, pH 7.4 and CP2-CP1 supplemented with 0.5 % w/v fatty acid-free bovine serum albumin, 2 mM MgCl_2,_ 1 mM ATP and 250 U•0.1ml^-1^ subtilisin protease type VIII, pH 7.4] as described by (60). Heart mitochondria were isolated from 1-2 wildtype and Tazkd mice. Mice were euthanized by cervical dislocation and the heart was immediately excised, excess blood was removed, and the heart was placed in ice-cold buffer B [0.25 M sucrose, 10 mM HEPES (pH 7.2) and 5 mM EGTA]. After few seconds to allow the remaining blood to decant, the heart was rapidly diced into very small pieces and the suspension was poured into a 30 mL ice-jacket Potter-Elvehjem with motorized Teflon pestle and 10-15 mL buffer A [buffer B supplemented with 0.5 % w/v fatty acid-free bovine serum albumin] was added. After 6-10 strokes the homogenate was transferred to an ice-cold 50 mL centrifuge tube and spun at 500 x g for 5 min. The supernatant was transferred to a clean tube and spun at 10,000 x g for 10 min. Now, the supernatant was discarded, and the pellet was carefully resuspended in 0.5 mL buffer B. and then 20 mL buffer B was added followed by a quick centrifugation at 1000 x g for 3 min. The supernatant was transferred to a fresh tube and centrifuged at 10,000 x g for 10 min. The pellet was resuspended in 200 µL of buffer B and protein concertation was determined using bicinchoninic acid assay.

### Western Blot Analysis

To determine tafazzin levels, 10 µg samples of skeletal muscle and heart isolated mitochondrial protein were boiled in Laemmli loading buffer under reducing conditions. Proteins were separated by 4 –12% NU-PAGE gradient gel using 1x MOPS buffer (Invitrogen) and transferred to a nitrocellulose membrane. Anti-tafazzin (1:1000 dilution) was a kind gift from Dr. S.M. Claypool (61) and recognizes a band just below 30kDa. Chemiluminescence was generated with SuperSignal West Pico (Thermo Scientific) and quantified with Image J software (National Institutes of Health).

### Mitochondrial Oxygen Consumption and Superoxide/H_2_O_2_ Production

Oxygen consumption rate (OCR) of freshly isolated heart and skeletal muscle mitochondria was monitored using an XF-24 extracellular flux analyzer (Seahorse Bioscience, Agilent). Briefly, 2 µg of mitochondrial protein was plated per well in KHE medium (120 mM KCl, 5 mM HEPES, 1 mM EGTA) supplemented with 0.3% w/v fatty acid-free bovine serum albumin, 1 mM MgCl_2_, 5 mM KH_2_PO_4_. Baseline rates with 1 mM ADP were measured for 15 min and 5 mM succinate + 4 µM rotenone were injected from port A. OCR at state 3 was monitored for 9 min and 1µg•ml^-1^ oligomycin was injected from port B to induce mitochondrial state 4.

Rates of superoxide/H_2_O_2_ production were collectively measured as rates of H_2_O_2_ generation, as two superoxide molecules are dismutated by mitochondrial or exogenous superoxide dismutase to yield one H_2_O_2_. The maximum capacities for superoxide/H_2_O_2_ production from the eleven sites were detected fluorometrically in a 96-well plate using specific combinations of inhibitors and substrates (see below) in the presence of 5 U•ml^-1^ horseradish peroxidase, 25 U•ml^-1^ superoxide dismutase and 50 µM Amplex UltraRed (56). Heart and skeletal muscle mitochondria (0.1 mg protein•ml^-1^) were incubated in KHE medium supplemented with 0.3% w/v fatty acid-free bovine serum albumin, 1 mM MgCl_2_, 5 mM KH_2_PO_4_, 1 µg•ml^-1^ oligomycin. Changes in fluorescence signal (Ex 604/Em 640) were monitored for 15 min. using a microplate reader (Molecular Devices SpectraMax Paradigm) and calibrated with known amounts of H_2_O_2_ at the end of each run in the presence of all inhibitors and effectors (56). The sites linked to the NADH/NAD^+^ isopotential pool were measured in the presence of 4 µM rotenone: Site I_F_: 5 mM malate, 2.5 mM ATP and 5 mM aspartate; site A_F_: 10 mM 2-oxoadipic acid; B_F_: 20 mM KMV (3-methyl-2-oxopentanoate/alpha-keto-methylvalerate); P_F_: 2.5 mM pyruvate and 5 mM carnitine; O_F_: 2.5 mM 2-oxoglutarate and 2.5 mM ADP. Site I_Q_ was measured as the rotenone-sensitive rate in the presence of 5 mM succinate. The remaining sites were linked to the ubiquinone isopotential pool: Site III_Qo_ was measured as the myxothiazol-sensitive rate in the presence of 5 mM succinate, 5 mM malonate, 4 µM rotenone and 2 µM antimycin A. Site II_F_ was measured as the 1mM malonate-sensitive rate in the presence of 0.2 mM succinate and 2 µM myxothiazol. Site G_Q_: 10 mM glycerol-3-phosphate, 4 µM rotenone, 2 µM myxothiazol, 2 µM antimycin A and 1 mM malonate; Site D_Q_: 3.5 mM dihydroorotate, 4 µM rotenone, 2 µM myxothiazol, 2 µM antimycin A and 1 mM malonate; Site E_F_: 15 µM palmitoylcarnitine, 2 mM carnitine, 5 µM FCCP, 1 mM malonate, 2 µM myxothyazol.

For the measurement of the rate of H_2_O_2_ production under native conditions *ex vivo*, skeletal muscle and heart mitochondria (0.1 mg of protein•ml^-1^) from 12 month-old wildtype and tazkd mice were incubated at 37 °C for 4 –5 min in the appropriate “basic medium” mimicking the cytosol of skeletal muscle or heart during rest, respectively, (basic “rest” medium plus 1µg•ml^-1^ oligomycin) and then added to a black 96 well plate containing the complex substrate mix. Changes in fluorescence signal (Ex 604/Em 640) were monitored for 15 min. using a microplate reader (Molecular Devices SpectraMax Paradigm) and calibrated with known amounts of H_2_O_2_ at the end of each run in the presence of all substrates.

### Skeletal muscle basic medium

40 mM taurine, 3 mM KH_2_PO_4_, 3.16 mM NaCl, 52.85 mM KCl, 5.46 mM MgCl_2_ (targeted free Mg^2+^ concentration 600 µM), 0.214 mM CaCl_2_ (targeted Ca^2+^ concentration 0.05 µM), 10 mM HEPES, 1 mM EGTA, 0.3 % w/v fatty acid-free bovine serum albumin, 1 µg•ml^-1^ oligomycin, pH 7.1. Targeted Na^+^ concentration was 16 mM and total K^+^ concentration was 80 mM. The medium had K^+^ and Cl^-^ adjusted to give an osmolarity of 290 mosM. Total Mg^2+^ and Ca^2+^ concentrations to give the targeted free values were calculated using the software MaxChelator (26).

### Skeletal muscle complex substrate mix

100 µM acetoacetate, 300 µM 3-hydroxybutyrate, 2500 µM alanine, 500 µM arginine, 1500 µM aspartate, 1500 µM glutamate, 6000 µM glutamine, 7000 µM glycine, 150 µM isoleucine, 200 µM leucine, 1250 µM lysine, 2000 µM serine, 300 µM valine, 100 µM citrate, 200 µM malate, 30 µM 2-oxoglutarate, 100 µM pyruvate, 200 µM succinate, 100 µM glycerol-3-phosphate, 50 µM dihydroxyacetone phosphate, 1000 µM carnitine, 500 µM acetylcarnitine, 10 µM palmitoylcarnitine and 6000 µM ATP (26).

### Heart basic medium

70 mM taurine, 6 mM KH_2_PO_4_, 6 mM NaCl, 12.8 mM KCl, 9.2 mM MgCl_2_ (targeted free Mg^2+^ concentration 1200 µM), 0.42 mM CaCl_2_ (targeted Ca^2+^ concentration 0.1 µM), 10 mM HEPES, 1 mM EGTA, 0.3 % w/v fatty acid-free bovine serum albumin, 1 µg•ml^-1^ oligomycin, pH 7.1. Targeted Na^+^ concentration was 18 mM and total K^+^ concentration was 55.5 mM. The medium had K^+^ and Cl^-^ adjusted to give an osmolarity of 284 mosM. Total Mg^2+^ and Ca^2+^ concentrations to give the targeted free values were calculated using the software MaxChelator.

### Heart complex substrate mix

50 µM 3-hydroxybutyrate, 2000 µM alanine, 400 µM arginine, 4500 µM aspartate, 5000 µM glutamate, 8000 µM glutamine, 1000 µM glycine, 80 µM isoleucine, 130 µM leucine, 800 µM lysine, 350 µM Proline, 1000 µM serine, 150 µM valine, 100 µM citrate, 150 µM malate, 50 µM 2-oxoglutarate, 130 µM pyruvate, 135 µM succinate, 400 µM glycerol-3-phosphate, 40 µM dihydroxyacetone phosphate, 1700 µM carnitine, 80 µM acetylcarnitine, 5 µM palmitoylcarnitine and 9000 µM ATP.

### Isolation and quantification of cardiolipin (CL) and monolysocardiolipin (MLCL)

Lipids were extracted from isolated heart and skeletal muscle mitochondria using methanol chloroform (2:1). Isolation and quantification of cardiolipin and monolysocardiolipin were performed according to (62).

### Body composition, Comprehensive Lab Animal Monitoring System (CLAMS) and exercise challenge

Body composition from wildtype and Tazkd mice at 3 and 6 months of age was measured with dual-energy x-ray absorptiometry (DEXA Lunar PIXImus, GE). Animals were anesthetised with 100 mg/kg ketamine and 10 mg/kg xylazine. For CLAMS, mice were housed individually and acclimatized for 1 day. Oxygen consumption, carbon dioxide release, energy expenditure and activity were measured using a Columbus Instruments Oxymax-CLAMS system according to guidelines for measuring energy metabolism in mice (63, 64).

Mice were physically challenged on a lidded motorized treadmill (Columbus Instrument, Columbus, OH, USA), which had adjustable speed and inclination and electric shock stimulation grid. The stimulus intensity was set to 1 mA. Mice were acclimatized in the treadmill for 3 d before the test. Acclimation protocol: 5 m/min for 5 min, followed by 1 m/min increment in speed every min until 10 min. Mice were allowed to rest for 5 min and then run for 10 min at 10m/min. On the test day, wildtype and tazkd mice were placed in the treadmill and the test started with the mice running for 5 min at 5m/min, then the speed was increased 1 m/min every min up to 20 min. Exhaustion was achieved when a mouse stayed more than 10 s on the stimulus grid or touched the grid more than ten consecutive times.

### Statistics

Data are presented as mean ± SEM of n independent values. Student’s t-test was used to calculate statistical significance when two groups were compared (Figures 2, 3 and 6). For multiple comparison tests where age, genotype and sites of superoxide/hydrogen peroxide production were analyzed, 2-way ANOVA followed by Sidark’s post-hoc test was used and when appropriate Bonferroni correction was applied. P<0.05 was considered statistically significant (GraphPad Prism).

### Data availability

All the data presented here is contained in this manuscript.

## Acknowledgments

The authors thank members of the Hotamisligil laboratory, especially Dr. Karen Inouye for her valuable help in the mice facility and Dr. Günes Parlakgül for the comments. We are grateful to Dr. Steven Claypool for kindly providing an aliquot of anti-tafazzin antibody, and Michael MacArthur for assisting on the treadmill experiments.

## The abbreviations used are

H_2_O_2_: hydrogen peroxide
Tazkd: tafazzin knock down
BTHS: Barth Syndrome
FAD: flavin adenine mononucleotide
CL: cardiolipin
MLCL: monolysocardiolipin
ROS: reactive oxygen species
site I_F_: flavin in the NADH-oxidizing site of respiratory complex I
site I_Q_: ubiquinone-reducing site of respiratory complex I
site II_F_: flavin site of respiratory complex II
site III_Qo_: outer quinol-oxidizing site of respiratory complex III site A_F,_ flavin in 2-oxoadipate dehydrogenase
site O_F_: flavin in the 2-oxoglutarate dehydrogenase complex
site P_F_: flavin in the pyruvate dehydrogenase complex
site B_F_: flavin in the branched-chain 2-oxoacid (or *α*-ketoacid) dehydrogenase complex
site G_Q_: quinone reducing site in mitochondrial glycerol 3-phosphate dehydrogenase, mGPDH
site E_F_: site in the electron transferring flavoprotein/ETF:ubiquinone oxidoreductase (ETF:QOR), probably the flavin of ETF
site D_Q_: quinone reducing site in dihydroorotate dehydrogenase Q, ubiquinone
QH_2_: ubiquinol
E_h_: operating redox potential
RER: respiratory exchange ration
EE: energy expenditure
S1QEL: <u>s</u>uppressor of site <u>I_Q_ e</u>lectron leak
S3QEL: <u>s</u>uppressor of site <u>III</u>_<u>Q</u>o_ <u>e</u>lectron leak
mCAT: mitochondrial-targeted catalase

## Supplementary information includes

Supplementary Table 1

Supplementary Figure 1

**Supplementary figure 1.**
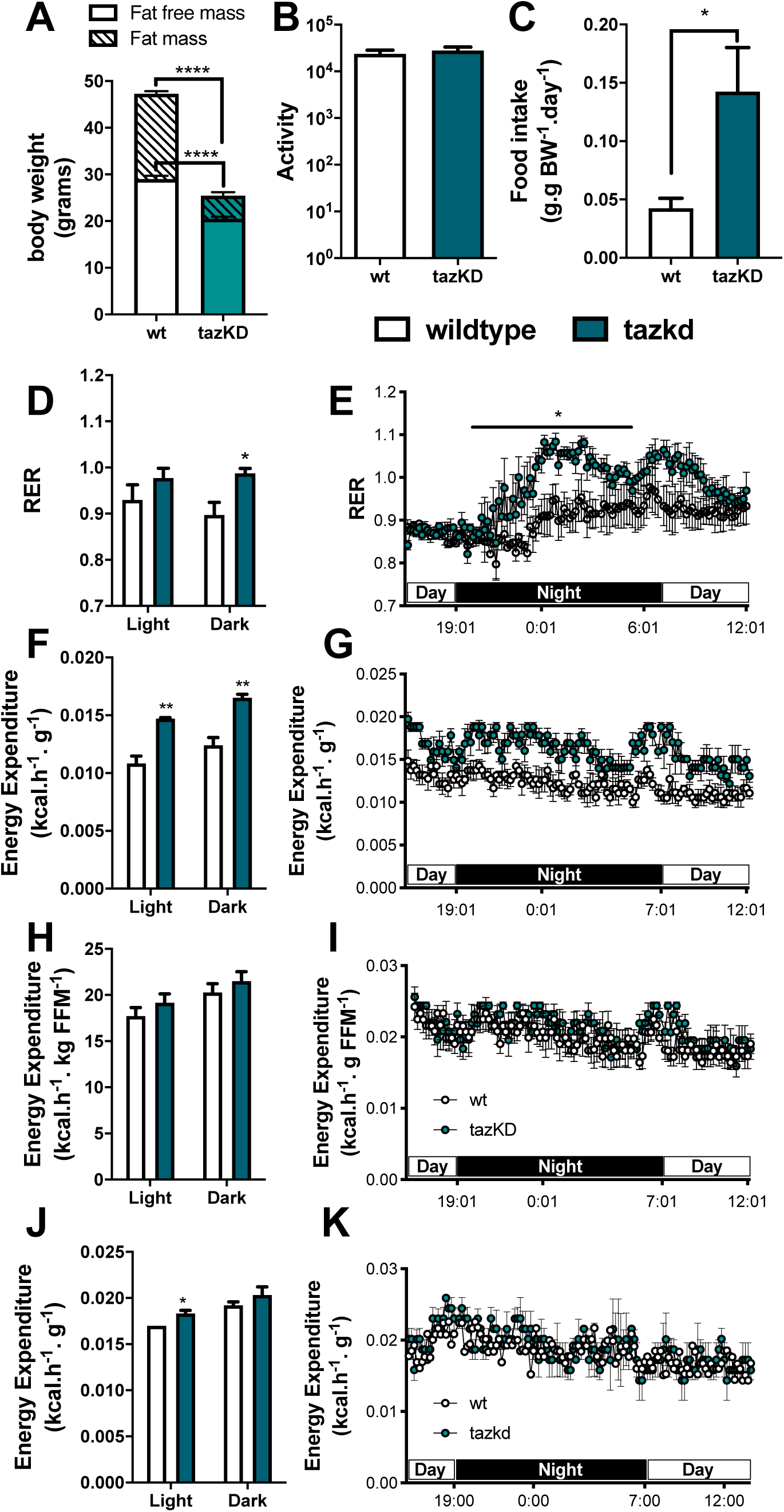
Metabolic profile of tazkd mice. **A**. body weight, fat mass and fat-free mass of 7-month old wildtype and tazkd mice. **B**. activity, measured by the number of beam breaks in the cage. **C**. food intake normalized by body weight. **D** and **E**, histogram and representative plot of respiratory exchange ratio (RER) of wildtype and tazkd mice over a period of 24 h. **F** and **G**, histogram and representative plot of energy expenditure (normalized by body weight) of wildtype and tazkd mice at 6 months of age over a period of 24 h. **H** and **I**, replot from F and G normalized by fat-free mass. **J** and **K**, histogram and representative plot of energy expenditure (normalized by whole body weight) of wildtype and tazkd mice at 3 months of age over a period of 24 h. Dark and light cycles are shown. Values are mean ± SEM, n=4. *, p<0.05, using Student’s t-test. ****, p<0.0001, 2-way ANOVA, Sidak’s post test.

**Supplementary Table 1.** Metabolite concentrations at rest in cytosol of heart and skeletal muscle and the composition of “ex vivo” media mimicking cytosol at rest. Data for the metabolites assumed relevant, from literature for rat heart (cited here) and skeletal muscle (from (1)). For heart, calculations assume heart wet weight is 79% water, 79% of which is intracellular (5, 29, 47–49). Values are corrected for distribution of intracellular metabolites between cytosol and mitochondria where known (24, 36, 50), otherwise assuming liver distribution (51–56), and that mitochondrial volume is 25% of intracellular (47, 48). Values are rounded to convenient whole numbers. Total Mg^2+^ and Ca^2+^ concentrations to give the targeted free values were calculated using the software MaxChelator. Targeted free calcium concentrations were 0.1 µM (heart) and 0.05 µM (skeletal muscle). Targeted free Mg^2+^ concentrations were 1200 µM (heart) and 600 µM (skeletal muscle). Targeted Na^+^ concentrations were 18 mM (heart) and 16 mM (skeletal muscle). Media had K^+^ and Cl^-^ adjusted as shown to give a target osmolarity of 290 mOsm (actual 284 mOsm) (57–59). Total K^+^ concentrations were 55.5 mM (heart) and 80 mM (skeletal muscle). Both media were supplemented with 0.3 g BSA/100 ml, 10 mM Hepes, 2 mM EGTA, 2 µg/mL oligomycin. “Rest” metabolite concentrations were mean literature values from the cited references for the hearts of animals at rest or from isolated Langendorff perfused heart beating at the spontaneous rate.

| Metabolite | Literature value for concentration in cytosol and value in medium mimicking cytosol ($\mu\text{M}$) | | References for heart metabolite concentrations (for skeletal muscle references see (1)) |
| --- | --- | --- | --- |
|  | Heart | Skeletal Muscle |  |
| Acetoacetate | 0 | 100 | (2) |
| 3-Hydroxybutyrate | 50 | 300 | (2) |
| Glycerol 3-phosphate | 400 | 100 | (2–7) |
| Dihydroxyacetone phosphate | 40 | 50 | (3, 5–9) |
| Citrate | 100 | 100 | (2, 3, 15–23, 4, 5, 9–14) |
| Malate | 150 | 200 | (2, 10–13, 16–18, 21, 23–25) |
| 2-oxoglutarate | 50 | 30 | (2, 10–12, 14–18, 21, 23–26) |
| Pyruvate | 130 | 100 | (2, 5–7, 9, 13, 17, 18, 24) |
| Succinate | 135 | 200 | (10, 15, 18, 25, 27) |
| Alanine | 2000 | 2500 | (11, 14, 15, 17, 18, 23, 28, 29) |
| Arginine | 400 | 500 | (15, 23, 28) |
| Aspartate | 4500 | 1500 | (7, 10–18, 21, 23, 28, 29) |
| Glutamate | 5000 | 1500 | (7, 10–18, 21, 23, 26, 28, 29) |
| Glutamine | 8000 | 6000 | (14, 15, 18, 23, 29) |
| Glycine | 1000 | 7000 | (14, 15, 23, 28, 29) |
| Isoleucine | 80 | 150 | (14, 23, 28–30) |
| Leucine | 130 | 200 | (14, 23, 28–30) |
| Lysine | 800 | 1250 | (15, 23, 28) |
| Proline | 350 | 500 | (28) |
| Serine | 1000 | 2000 | (14, 15, 23, 28, 29) |
| Taurine | 70000 | 40000 | (14, 15, 23, 28, 31) |
| Valine | 150 | 300 | (14, 28–30) |
| Carnitine | 1700 | 1000 | (32–36) |
| Acetylcarnitine | 80 | 500 | (12, 13, 33, 35) |
| Palmitoylcarnitine | 5 | 10 | (37) |
| Basic media |  |  |  |
| ATP | 9000 | 6000 | (2, 4, 5, 7, 12, 13, 20, 23, 26, 38–42) |
| $\text{KH}_2\text{PO}_4$ | 6000 | 3000 | (5, 7, 9, 23, 26, 38, 40, 42) |
| EGTA | 2000 | 2000 |  |
| KCl | 12790 | 52850 |  |
| NaCl | 6000 | 3167 | (43) |
| $\text{MgCl}_2$ | 9224 | 5467 | (9) |

## Footnotes

1 Colin Phoon et al., communication at the Barth Syndrome Foundation Conference in July 2018, Florida.

